# Multi-week digital home cage monitoring reduces noise and enhances reproducibility

**DOI:** 10.1101/2025.10.29.685181

**Authors:** Michael C. Saul, Natalie Bratcher-Petersen, Manuel E. Ruidiaz, Juan Pablo Oberhauser, Vivek M. Philip, Susan E. Bolin, Brianna N. Gaskill, Timothy L. Robertson

## Abstract

Reproducibility is a persistent challenge in preclinical research. We used multi-week rodent machine vision home cage monitoring at three different pharmaceutical companies to examine factors governing replication of genotype differences in activity. Interlaboratory replication of genotype effects was surprisingly high. Longer study durations reduced noise, improving replication and reducing replicable sample sizes. These findings demonstrate the potential of long-term home cage digital monitoring as a method to enhance reproducibility.

## Introduction

Results that fail to reproduce pose a challenge in preclinical research. Over-standardization, improper or underpowered study designs, uncaptured environmental variability, experimenter influences on subjects, and insufficient protocol specification and harmonization can cause irreproducible research (Karp et al., 2020; Nigri et al., 2022; Richter et al., 2009). Paradigms that are not automated, performed in irregular environments, use short-duration methods, and occur during hours mice would typically be asleep further complicate reproducibility (Crabbe et al., 1999; Von Kortzfleisch et al., 2022). Further, unmeasured environmental factors and interlaboratory variation complicate reproducibility in ideal experiments (Crabbe et al., 1999; Von Kortzfleisch et al., 2022).

Enhancing reproducibility in preclinical research has great translational value. Suggested methods for enhancing reproducibility include reassessing species selection, utilizing systematic heterogenization, performing studies at different times of day, capturing cage effects, and controlling for factors like weaning time and housing (Bailoo et al., 2020; Bodden et al., 2019).

One potential method to improve preclinical research reproducibility is digital home cage monitoring (Pernold et al., 2019). Continuous long-duration measurement of behavior and physiology in the animal’s undisturbed home environment minimizes variability and enables assessment of the impact of human interaction. The Digital In Vivo Alliance (DIVA), a collaboration led by The Jackson Laboratory, aims to develop and validate digital measures and enhance translational and preclinical research. The JAX Envision™ platform facilitates scalable real-time monitoring of group-housed individuals in home cage environments. DIVA has used Envision™ for detailed protocol harmonization, automated continuous measurement, unbiased operator-independent assessments, and to reduce human–animal interaction.

We sought to understand how variation across three member sites contribute to measured biological effects in home cage activity. Unbiased digital measures collected in the home cage allow extended duration data collection measured in weeks. We posited that digital home cage phenotyping would improve inter-site replication through enhanced capture of sources of variation within and across sites.

## Brief Methods

We recorded home cage activity of both sexes of three mouse genotypes (C57BL/6J, A/J, and J:ARC) in three different vivaria (AbbVie in Lake County, IL; Novartis in San Diego, CA; and BioMarin in Marin County, CA). Mice were housed in a 14L:10D light/dark cycle with lights-on at 04:00 local time and lights-off at 18:00 local time for all sites. All mice shipped from the Ellsworth, Maine facility of The Jackson Laboratory on three ship dates. Sites agreed on a study protocol and standardized as many variables as possible (light cycle, bedding, lot/batch of rodent diet, enrichment, cage change schedule, and handling practices; see: **Supplementary Methods**).

Mice were housed in groups of three, acclimated for three days, then tagged with custom RapID® tags (RapID Labs, San Francisco, CA) for individual animal identification. Mice were tracked for a further 18 days. We utilized an ANOVA model from previously published work (Crabbe et al., 1999):

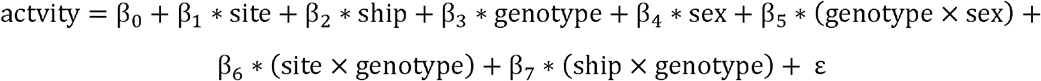

Genotype, sex, and their interaction were considered biological factors of interest; site, ship date, and their interactions with genotype were considered technical/environmental factors.

## Results and Discussion

We tracked a total of 76,495 hours (8.73 years) of mouse home cage activity across the 3 sites. During the first ship date, mice two sites experienced dehydration, resulting in early study termination for one cages and addition of Hydro-Gel® (Clear H2O, Westbrook, ME) for this and subsequent trials. During the final ship date, multiple days of video data were lost due to an error.

The experiment was designed with the cage as the experimental unit and data were analyzed with cage-level activity as the primary dependent variable. There was substantial variation in average activity within each cage as measured by 24-hour rolling means (**Figure 1A**). Ship date explained 31.7% of the variance in the hourly coefficient of variation for detrended data (F_2,36_ = 16.47, p < 0.001, **Figure 1B**). The higher variation in the earliest ship date may reflect noise introduced by the adverse husbandry events noted above.

**Figure 1.**
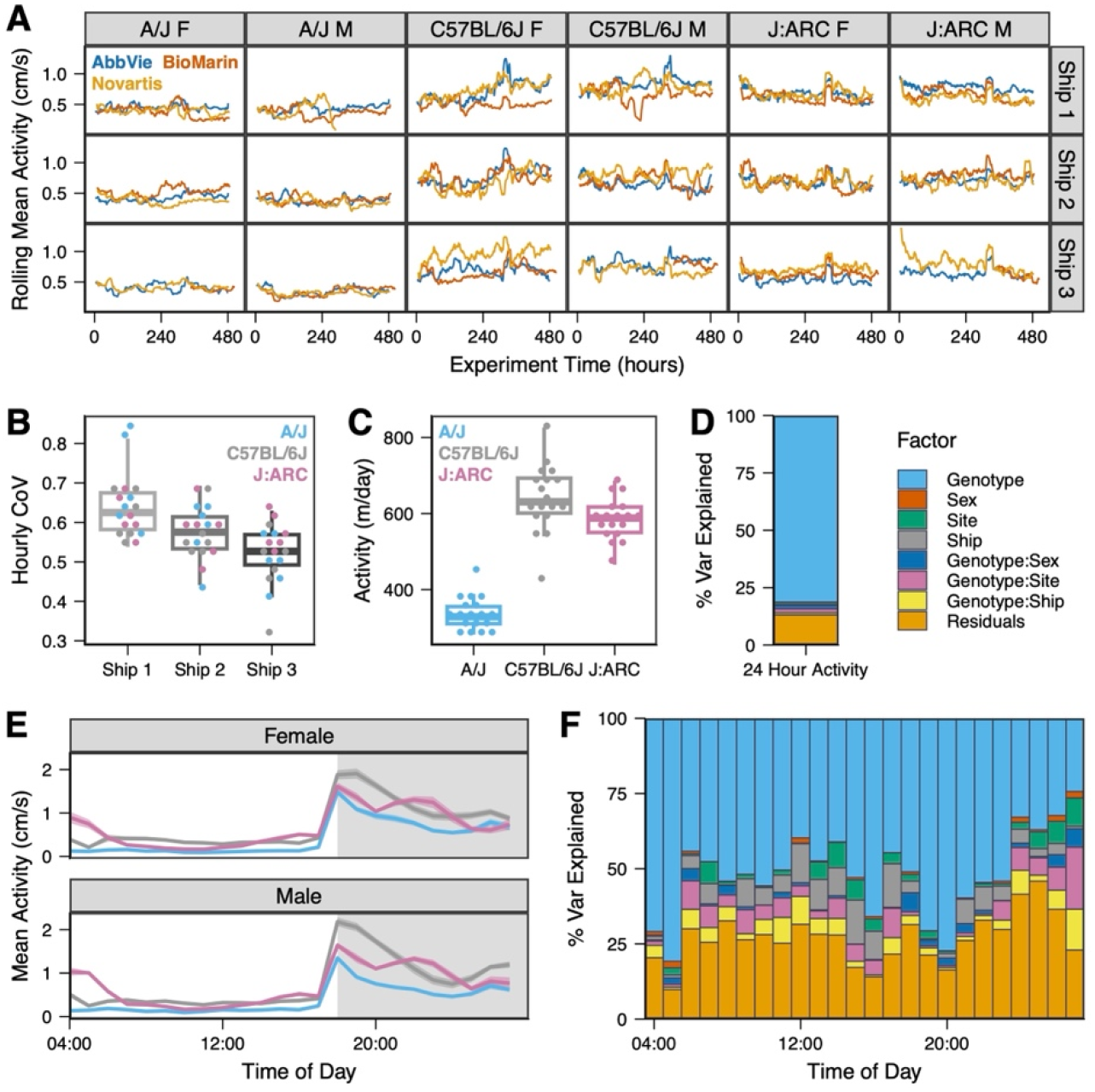
Genotype effect scales with duration. **A)** Variation in 24-hour moving average data. **B)** Detrended CoV differs across ship dates. **C)** Genotype strongly governs daily activity at full study duration. **D)** Genotype explains over 80% of the variance in daily activity. **E)** Genotypes have characteristic highly stereotyped circadian patterns. **F)** Genotype effect strength depends on time of day.

When the data were taken in aggregate, genotype explained 81.3% of the variance in average 24-hour activity (**Figure 1C-D**). Technical factors explained 3.9% of the variance and residuals explained 13.1% of the variance in average activity, indicating that both technical factors and noise were minimal in the aggregate measures. Sex and its interaction with genotype explained the final 1.7% of the variance. Further, each site independently identified strong genotype effects on 24-hour activity (75.5%-88.7% variance explained, AbbVie: F_2,6_ = 36.24, p < 0.001; BioMarin: F_2,6_ = 39.82, p < 0.001; Novartis: F_2,6_ = 36.11, p < 0.001), demonstrating replication between these sites.

Data were further averaged by time of day. Each genetic background and sex displayed its own characteristic circadian pattern (**Figure 1E**). Each genotype was active in the dark period and inactive in the light period. Consistent with previous measurement (Logan et al., 2013), C57BL/6J generally showed the highest activity, J:ARC was intermediate, and A/J was lowest. J:ARC activity remained relatively stable across the dark period. Genotype effect strength depended upon time of day (**Figure 1F**); genotype effects dominated at specific times (04:00- 05:00, 18:00-23:00), while residuals (00:00-01:00), and technical factors (06:00-17:00, 02:00- 03:00) were stronger at other times. Thus, time of measurement influences site-to-site reproducibility of results.

To simulate how the duration of a study influences replication, each hour was randomly subsampled 100 times to between 1 and 20 days (**Supplementary Methods**). Each subsample yielded approximately the same sample size, allowing direct comparison between experiment durations.

Shorter duration was generally associated with greater experimental noise. As subsample increased, rescaled variance explained by technical and biological factors also increased (**Figure 2A**). ANOVAs for each site demonstrate that replication as measured by simultaneous significant results between sites increased for subsamples with longer durations (**Figure 2B**). In 20 day subsamples, genotype effect replication varied by the time of day when tests were performed (**Figure 2C**). These results indicate that reproducibility is enhanced both by longer study duration and by using 24-hour continuous monitoring to capture replicable times when experimenters would typically be absent.

**Figure 2.**
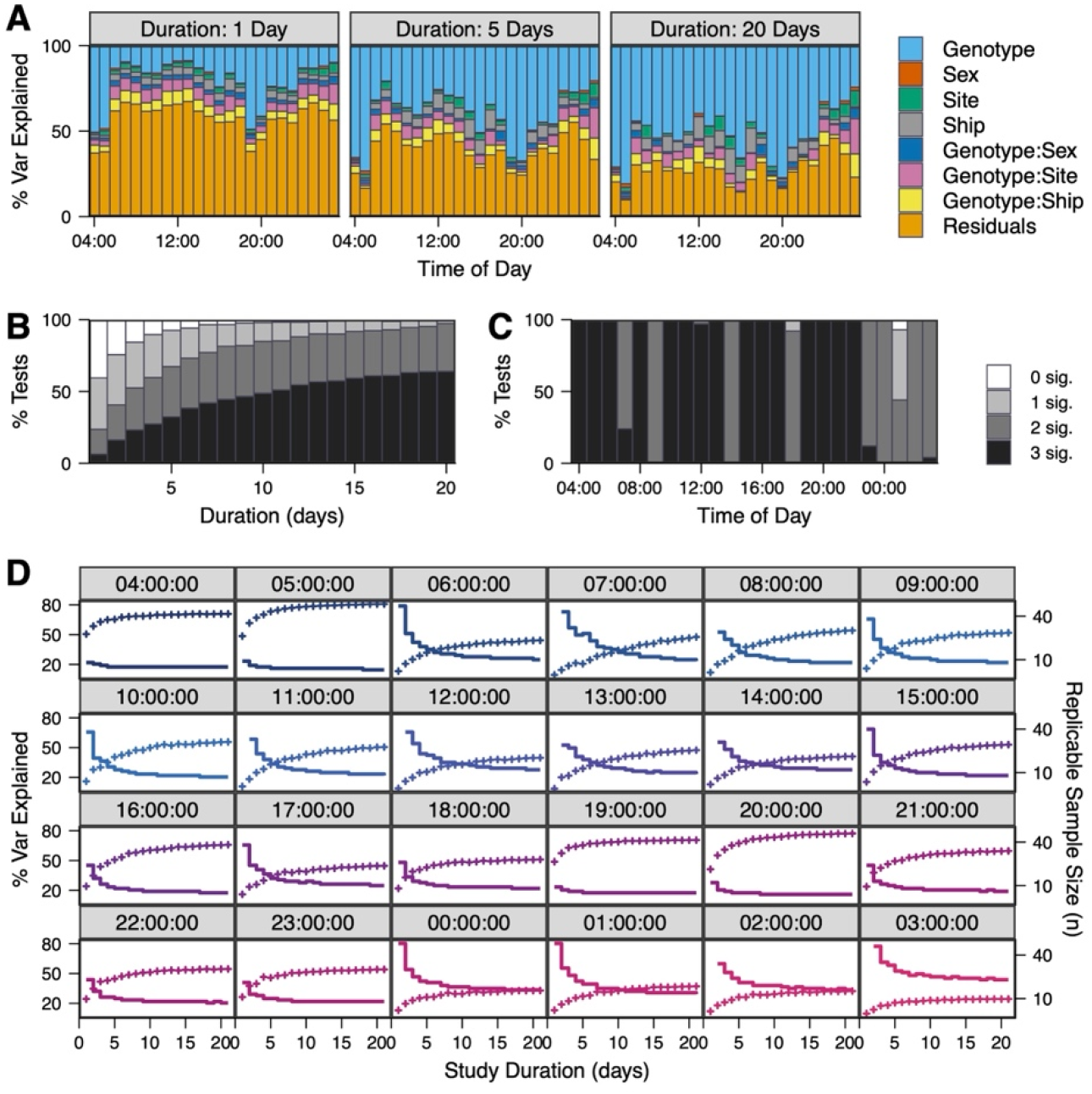
Results of hourly subsamples. **A)** Residual variance decreases with longer subsample duration. **B)** Longer subsamples increase likelihood of genotype effect replication. Black indicates replication at all three sites. **C)** Multi-site replication with 20-day subsamples depends upon time of day. **D)** Percent variance explained (crosses) increases with subsample duration. Sample size needed for well-powered replicable experiments (solid stair-stepped line) decreases as duration increases.

Subsample effect sizes were used to perform power analysis for sample sizes needed to replicate across three sites (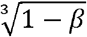 where *β*= 0.2 using one-way ANOVAs on three classes for *p* = 0.05). For a single day duration, the median number of cages needed for replicable results was n=39 with a range between n=8 at 04:00 to n=75 at 13:00. Sample sizes for single day durations appeared relatively high between 10:00 and 15:00, (**Figure 2D**), when workers are often present and when data is often collected. For 20 days’ duration, the median sample size for replicable results reduced to n=8 and ranged from n=3 at 05:00 to n=23 and 03:00 (**Figure 2D**). The results of the power analysis indicate that short duration testing for 1 hour or less during the time when experiments are typically performed requires very high sample sizes to achieve replicable results, but that sample sizes may be reduced both by testing at different circadian times and by averaging over many days.

Long-duration monitoring allowed us to overcome substantial day-to-day noise and increase inter-laboratory replication. Multi-week designs with many data points per biological replicate averaged together enhance reproducibility. Further, reproducibility appears to be improved through unbiased measurement of behavioral and physiological endpoints during hours when animals are typically awake and humans are not present in the animal room. We note that these findings suggest that increased variability in the light period resulted from circadian influences or from the presence of humans in the room at these times, but that the present dataset cannot adjudicate this hypothesis. Future work on reproducibility would benefit from examining such additional sources of variability that we were unable to assess here.

## Acknowledgments

We than Kristin Evans and Brian Baridon from BioMarin, Christina Boykin and Joseph Brewster from Novartis, the AbbVie research training services, and the three site animal husbandry teams. We thank Brian Berridge of B2 Pathology Solutions and DIVA chair for his thoughtful feedback. We acknowledge the Jackson Laboratory for providing the mice and the beta version of the technology used in data collection and DIVA for funding the work.

