## Supplementary Methods for "Multi-week digital home cage monitoring reduces noise and enhances reproducibility"

### Supplementary Materials and Methods

#### Animals

##### Experimental Animals

This study utilized animals from three different genetic backgrounds: the inbred C57BL/6J strain (JAX stock #000664, RRID:IMSR_JAX:000664), the inbred A/J strain (JAX stock #000646, RRID:IMSR_JAX:000646), and the outbred stock J:ARC(S) (JAX stock #034608, RRID:IMSR_JAX:034608). Males and females of each genetic background aged six weeks were taken from single housing rooms at the Ellsworth, ME facility of The Jackson Laboratory and shipped to the respective sites for this study.

##### Ethics Statement

All animal studies were conducted under an approved Institutional Animal Care and Use Committee (IACUC) protocol at each study site.

##### Study Design and Justification of Animal Numbers

Our experimental design was factorial using three locations, both males and females from three genetic backgrounds, and three experimental replicates derived from different ship dates for a total of 54 cages and 162 mice. Though each cage housed three mice, animals within a cage are confounded with the social milieu of the cage itself. Thus, the cage was considered the experimental unit. Mead’s Resource Equation was utilized during experimental design to ensure adequate replication of each genetic background and sex combination at each site. Though the resource equation indicated that two ship date replicates per genetic background and sex combination would have been sufficient for most within subjects' designs made possible by long-duration tracking, we noted in advance that any cage attrition would compromise the experimental design. Consequently, we elected to add a third ship date in advance to make the experimental design more robust against potential unforeseen issues.

##### Environment, Housing, Husbandry, and Experimental Apparatus

The enabling technology behind this study is a novel home cage tracking apparatus. Briefly, the apparatus used here is a beta version of the Allentown Discovery™ rack powered by the JAX Envision™ platform. The rack-based format of this beta version includes nine total cages in a rat form factor. Cage bottoms were commercially available disposable rat cages (Innovive, San Diego, CA, USA) with customized cage lids that permitted a top-down view. Each individual home cage included both visible and far-infrared LED lights to enable continuous monitoring during both the light and dark cycles as well as a top-down camera to capture all motion within each cage. The racks continuously collected video and uploaded the data to the Amazon cloud, where the data were analyzed using a prototype of the JAX Envision™ platform.

Mice were trio housed in these home cages on irradiated aspen chip bedding (Sani-Chips®, P.J. Murphy Forest Products Corporation, Montville, NJ) with 4g of brown shredded paper (Bed’r Nest®, The Andersons Plant Nutrients, Maumee, OH, USA) and 2 cotton square nestlets (Ancare Corp, Bellmore, NY) for enrichment. Mice were provided with reverse osmosis chlorinated water acidified to pH 2.4 - 2.8 and delivered via bottle with surface tension tube. Mice were fed ad libitum with Teklad Global 2014 14% Protein Rodent Maintenance Diet (Envigo, Madison, WI, USA). Animals were kept in rooms with a14:10 light/dark cycle (lights-on at 04:00, lights-off at 18:00) within each facility. Rooms were kept at a set point temperature of 22° C +/- 1° C.

##### Animal Handling

As much as possible, animal handling was standardized between the sites. During cage change, some of the existing nest was transferred to the new cage. Mice were handled manually by grasping the base of the tail using hands and not forceps. When possible, males were handled before females, and gloves were changed between handling of each sex.

#### Experimental Procedures

##### Blinding

Given the obvious visual differences between animals within the experiment and the impossibility of blinding to location and temporal replicate, blinding was not considered during the conduct of the experiment. The experimental outcome measures were collected automatically while the mice were housed in their cages. A factorial design with no *a priori* directional hypotheses was planned. Additionally, analysis using this novel technology frequently required an analyst to visually assess videos of animals, which would unblind the analyst to experimental conditions. Consequently, a blinded analysis was not considered.

##### Randomization

Within each site, home cages were assigned to random locations in their racks in advance. The randomization strategy was balanced to ensure equal sampling of each strain and each sex within each row and column of the racks.

##### Animal identification

Animals were lightly anesthetized with an inhalant anesthetic (Isoflurane, 2-3%) in an induction chamber until loss of righting reflex and then transferred to a heated platform and maintained on isoflurane anesthesia (2%) using a nose cone. Custom RapID ear tags designed for computer vision identification were placed in both ears with each tag facing opposite directions. Ear tags included a red tag with a black and white vertical stripe, a brown tag with a checkerboard pattern, and a black tag with an all-white body. Upon recovery from ear tagging, animals were returned to their home cages.

##### Continuous home cage monitoring

Animals were observed for a total study duration of 14-days. This observation included the following phases:

| Event/Phase | Study Day | Time/Duration | Description |
| --- | --- | --- | --- |
| Facility quarantine/acclimation | -14 to -7 | 7 days | Acclimation to facility/quarantine period for 7 days in Innovive caging in the DAX cage system environment. Recording to begin at arrival to capture acclimation to facility following transport. |
| Ear tag placement | -11 |  | 72 hours after unboxing (+/- 2 hours) |
| Cage change | 0 | 6-8 am | 2-4 hours from lights on |
| Home cage monitoring | 0 to 7 | 7 days | Continuous monitoring of mice in their home cage |

Upon arrival, animals were placed directly into DAX cages (rack-based). Seventy-two hours (+/- 2 hours) after unboxing and placing into DAX cages, animals were lightly anesthetized for ear tag placement (Animal identification section) and placed back into the DAX cages.

##### Outcome Measures

The primary outcome measure utilized in this study was cage-level activity as measured by activity over time in the home cage. This measure is derived from continuous video monitoring based upon the computed movement of mouse tracked by Euclidean distance from one frame to another. This outcome measure is expressed in centimeters per second (cm/s) and can be measured at a time scale as granular a resolution as 1 second. In this study, the cm/s traveled by animals each hour normalized to the number of animals per cage is the primary endpoint. An additional endpoint of 24-hour activity normalized to number of animals per cage as measured by meters moved in 24 hours is reported.

#### Analysis

##### Machine Vision Analysis

Video collected was analyzed using a prototype of the JAX Envision™ platform. Briefly, this system uses a set of machine vision algorithms to segment out the mice in the cage followed by a separate set of algorithms to perform individual tracking. Briefly, the videos collected by the cameras in the cage system were uploaded to AWS. These uploaded videos were then segmented for mice using a fast and efficient mHydra machine vision algorithm. Segmentation masks were converted into tracks with corresponding identities using a separate set of ID and tracking algorithms. The results of this activity suite were used to compute cage-level activity. A detailed technical description of the machine vision algorithms is provided in previous work^12^.

##### Statistical Analysis

All analysis was done in an Ubuntu 22.04.3 LTS x86_64 container on the Envision Developer’s Environment, an instance of the KubeFlow platform for machine learning. Analysis was performed on data exported from the Amazon S3 cloud. Analysis notebooks were written in the Quarto framework and all analysis was performed on R v4.3.1 and Python v3.9.

To facilitate cross-replicate and cross-site visualization and comparison, all data were aligned to the first transition to dark cycle after the animals were initially placed in the cage as T_0_.

For each pseudo experiment, we randomly selected a set of days without replacement from each hour within each cage to simulate capturing an arbitrary hour of an animal’s life. We then averaged the values together by hour and ran the same ANOVA model as above. For each duration, we performed 100 pseudo experiments, took the median percent variance explained for each factor, and rescaled these median values to add up to 100% to account for medians coming from different pseudo experiments.

Embargoed data used to produce the figures and analyses from this manuscript are posted to Figshare (private link URL: https://figshare.com/s/8ea317fbe134128314d9) and details of the statistical analysis are posted to GitHub (URL: https://github.com/msaul/diva-reproducibility-study-2025).
